# Cross-species alcohol dependence-associated gene networks: Co-analysis of mouse brain gene expression and human genome-wide association data

**DOI:** 10.1101/380584

**Authors:** Kristin M. Mignogna, Silviu A. Bacanu, Brien P. Riley, Aaron R. Wolen, Michael F. Miles

## Abstract

Genome-wide association studies on alcohol dependence, by themselves, have yet to account for the estimated heritability of the disorder and provide incomplete mechanistic understanding of this complex trait. Integrating brain ethanol-responsive gene expression networks from model organisms with human genetic data on alcohol dependence could aid in identifying dependence-associated genes and functional networks in which they are involved. This study used a modification of the Edge-Weighted Dense Module Searching for genome-wide association studies (EW-dmGWAS) approach to co-analyze whole-genome gene expression data from ethanol-exposed mouse brain tissue, human protein-protein interaction databases and alcohol dependence-related genome-wide association studies. Results revealed novel ethanol-regulated and alcohol dependence-associated gene networks in prefrontal cortex, nucleus accumbens, and ventral tegmental area. Three of these networks were overrepresented with genome-wide association signals from an independent dataset. These networks were significantly overrepresented for gene ontology categories involving several mechanisms, including actin filament-based activity, transcript regulation, Wnt and Syndecan-mediated signaling, and ubiquitination. Together, these studies provide novel insight for brain mechanisms contributing to alcohol dependence.

## Introduction

Alcohol Use Disorder [1], which spans the spectrum from abusive drinking to full alcohol dependence (AD), has a lifetime prevalence of 29.1% among adults in the United States [2]. Alcohol misuse ranks third in preventable causes of death in the U.S. [3] and fifth in risk factors for premature death and disability, globally [4]. Although pharmacological therapy for AUD exists [5], the effectiveness is limited and the relapse rate is high. Improvement in AUD treatment requires research on the underlying genetic and biological mechanisms of the progression from initial exposure to misuse, and finally to dependence.

Twin studies estimate that AUD is roughly 50% heritable [6, 7]. Multiple rodent model studies have used selective breeding to enrich for ethanol behavioral phenotypes or have identified ethanol-related behavioral quantitative trait loci [8–10], further confirming the large genetic contribution to alcohol behaviors. Recent studies have also documented genetic factors influencing the effectiveness of existing pharmacological treatments for AD, further substantiating genetic contributions to the mechanisms and treatment of AUD [11]. Genome-wide association studies (GWAS) in humans have identified several genetic variants associated with alcohol use and dependence [12–15]. However, they have yet to account for a large portion of the heritability estimated by twin studies. Lack of power, due to a large number of variants with small effects, is believed to the source of this “missing heritability”” [16]. Although recent large-scale studies have shown promise in identifying novel genetic contributions to alcohol consumption, these studies do not contain the deep phenotypic information necessary for identifying variants associated with dependence. Further, such GWAS results still generally lack information about how detected single gene variants are mechanistically related to the disease phenotype.

Genome-wide gene expression studies are capable of improving the power of GWAS by providing information about the gene networks in which GWAS variants function [17–20]. Although gene expression in brain tissue has been studied in AD humans [17, 18], these studies are often difficult to conduct and interpret, due to lack of control over experimental variables and small sample sizes. However, extensive studies in rodent models have successfully identified ethanol-associated gene expression differences and gene networks in brain tissue [21–24]. Multiple ethanol-behavioral rodent models exist to measure different aspects of the developmental trajectory from initial exposure to compulsive consumption [25]. Acute administration to naïve mice models the response of initial alcohol exposure in humans, which is an important predictor of risk for AD [26, 27]. Wolen et al. used microarray analysis across a mouse genetic panel to identify expression correlation-based networks of acute ethanol-regulated genes, along with significantly associated expression quantitative trait loci in the prefrontal cortex (PFC), nucleus accumbens (NAc), and ventral tegmental area (VTA) [24]. Furthermore, specific networks also correlated with other ethanol behavioral data derived from the same mouse genetic panel (BXD recombinant inbred lines) [10]. These results suggested that studying acute ethanol-exposed rodent brain gene expression could provide insight into relevant mechanistic frameworks and pathways underlying ethanol behaviors.

Several studies have integrated GWAS and gene expression or gene network data to cross-validate behavioral genetic finding [17]. For instance, the Psychiatric Genomics Consortium [28] tested for enrichment of nominally significant genes from human GWAS in previously identified functional pathways, and found shared functional enrichment of signals for schizophrenia, major depression disorder, and bipolar disorder in several categories. These pathways included histone methylation, neural signaling, and immune pathways [28]. Mamdani et al. reversed this type of analysis by testing for significant enrichment of previously identified GWAS signals in gene networks from their study. They found that expression quantitative trait loci for AD-associated gene expression networks in human prefrontal cortex tissue had significant enrichment with AD diagnosis and symptom count GWAS signals from the Collaborative Study on the Genetics of Alcoholism dataset [17]. Additional approaches have taken human GWAS significant (or suggestive) results for AD and provided additional confirmation by showing that expression levels for such genes showed correlations with ethanol behaviors in rodent models [29]. Such methods are informative with respect to analyzing the function of genes that have already reached some association significance threshold. However, they do not provide information about genes not reaching such statistical thresholds, but possibly still having important contributions to the genetic risk and mechanisms of AUD

Dense module searching for GWAS (dmGWAS) is an algorithm for directly integrating GWAS data and other biological network information so as to identify gene networks contributing to a genetic disorder, even if few of the individual network genes exceed genome-wide statistical association thresholds [30]. The initial description of this approach utilized Protein-Protein Interaction (PPI) network data to identify networks associated with a GWAS phenotype. Modules derived from protein-protein interactions were scored from node-weights based on gene-level GWAS *p*-values. This approach was used to identify AD-associated PPI networks that replicated across ethnicities and showed significant aggregate AD-association in independent GWAS datasets [31], thus demonstrating the potential utility of the method. A more recent iteration of the dmGWAS algorithm, termed Edge-Weighted dense module searching for GWAS (EW-dmGWAS), allows integration of gene expression data to provide a direct co-analysis of gene expression, PPI, and GWAS data [32].

Utilization of the EW-dmGWAS algorithm would allow for identification of gene networks coordinately weighted for GWAS significance for AD in humans and ethanol-responsiveness in model organism brain gene expression data. We hypothesized that such an approach could provide novel information about gene networks contributing to the risk for AUD, while also adding mechanistic information about the role of such networks in ethanol behaviors. We show here the first use of such an approach for the integration of human PPI connectivity with mouse brain expression responses to acute ethanol and human GWAS results on AD. Our design incorporated the genome-wide microarray expression dataset derived from the acute ethanol-exposed mouse brain tissue used in Wolen et al. [10, 24], human protein-protein interaction data from the Protein Interaction Network database, and AD GWAS summary statistics from the Irish Affected Sib-Pair Study of Alcohol Dependence [29]. Importantly, we validated the identified ethanol-regulated and AD-associated networks by co-analysis with an additional, independent AD GWAS study on the Avon Longitudinal Study of Parents and Children dataset. Our results could provide important methodological and biological function insight for further studies on the mechanisms and treatment of AUD.

## Materials and methods

### Sample

#### Mouse gene expression data

All mouse brain microarray data (Affymetrix GeneChip Mouse Genome 430 2.0) are from Wolen et al., 2012 [24] and can be downloaded from the GeneNetwork resource (www.genenetwork.org), via accession numbers GN135-137, GN154-156 and GN228-230, respectively for PFC, NAc and VTA data. Additionally, PFC microarray data is available from the Gene Expression Omnibus (GEO) via accession number GSE28515. Treatment and control groups each contained one mouse from each strain and were given IP injections of saline or 1.8 g/kg of ethanol, respectively. Euthanasia and brain tissue collection took place 4 hours later. Data used for edge weighting in EW-dmGWAS analysis included Robust Multi-array Average (RMA) values, background-corrected and normalized measures of probe-wise expression, from the PFC, VTA, and NAc of male mice in 27-35 BXD recombinant inbred strains and two progenitor strains (DBA/2J and C57BL/6J). For filtering of the same microarray datasets prior to EW-dmGWAS analysis (see below), we used probe-level expression differences between control and treatment groups determined in Wolen study using the S-score algorithm [33] (Table S1). Fisher’s Combined Test determined S-score significance values for ethanol regulation of each probeset across the entire BXD panel, and empirical p-values were calculated by 1,000 random permuations. Finally, q-values were calculated from empirical p-values to correct for multiple testing.

Ethanol-responsive genes are predicted to be involved in pathways of neural adaptations that lead to dependence [24]. We predicted they would also be involved in mechanistic pathways from which GWAS signals are being detected. We therefore performed a low-stringency filter for ethanol-responsiveness prior to EW-dmGWAS so as to ensure edge weighting focused on ethanol responsivity. To identify genes with suggestive ethanol responsiveness, we used a S-score probeset-level threshold of *qFDR*<0.1 for differential expression, in any one of the three brain regions. Genes associated with these probesets were carried forward in our analysis. Multiple probesets from single genes were reduced to single gene-wise expression levels within a particular brain region by selecting the maximum brain region-specific RMA value for each gene. After removing genes that were absent from the human datasets, 6,050 genes remained with expression values across all three brain regions (Fig 1).

**Fig 1.**
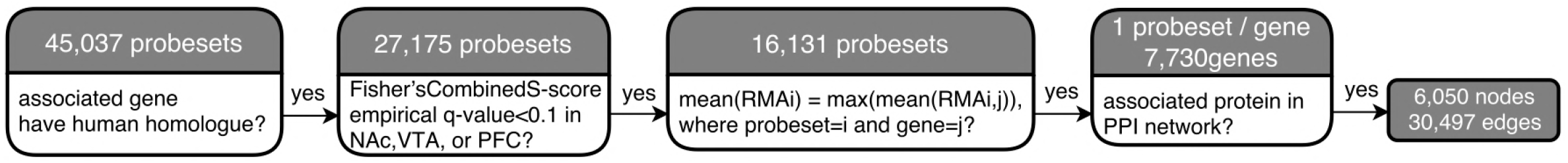
Data Pipeline for Determining Ethanol-Regulation and Merging Datasets. Pipeline used to prepare the data for the present analysis. The first cell contains the starting number of genes in the BXD mouse PFC, NAc, and VTA gene expression dataset.

#### Human GWAS data

The Irish Affected Sib-Pair Study of Alcohol Dependence (IASPSAD) AD GWAS dataset was used for the EW-dmGWAS analysis. It contains information from 1,748 unscreened controls (43.2% male) and 706 probands and affected siblings (65.7% male) from a native Irish population, after quality control [29]. Samples were genotyped on Affymetrix v6.0 SNP arrays. Diagnostic criteria for AD were based on the DSM-IV, and probands were ascertained from in- and out-patient alcoholism treatment facilities. Association of each Single Nucleotide Polymorphisms (SNP) with AD diagnosis status was tested by the Modified Quasi-Likelihood Score method [34], which accounts for participant relatedness. SNPs were imputed using IMPUTE2 [35] to hg19/1000 Genomes, and gene-wise p-values were calculated using Knowledge-Based mining system for Genome-wide Genetic studies (KGG2.5) [36].

The Avon Longitudinal Study of Parents and Children (ALSPAC) GWAS gene-wise p-values were used to examine the ability of EW-dmGWAS to validate the EW-dmGWAS networks. This GWAS tested SNP association with a factor score calculated from 10 Alcohol Use Disorder Identification Test items for 4,304 (42.9% male) participants from Avon, UK. Samples were genotyped by the Illumina HumanHap550 quad genome-wide SNP platform [37].

Although the analyzed phenotype was not identical to that in the IASPSAD GWAS, this dataset was similar to IASPSAD in that: 100% of the sample was European; the male to female ratio was roughly 1:1; SNPs were imputed to hg19/1000 Genomes; and gene-wise p-values were calculated by KGG2.5.

#### Protein network data

The Protein-Protein Interaction (PPI) network was obtained from the Protein Interaction Network Analysis (PINA 2.0) Platform (http://omics.bjcancer.org/pina/interactome.pina4ms.do). This platform includes PPI data from several different databases, including: Intact, MINT, BioGRID, DIP, HPRD, and MIPS/Mpact. The *Homo sapiens* dataset was used for this analysis [38, 39]. Uniprot IDs were used to match protein symbols to their corresponding gene symbols [40].

### Statistical methods

#### EW-dmGWAS

The edge-weighted dense module searching for GWAS (dmGWAS_3.0) R package was used to identify treatment-dependent modules (small, constituent networks) nested within a background PPI network (https://bioinfo.uth.edu/dmGWAS/). We used the PPI framework for the background network, IASPSAD GWAS gene-wise p-values for the node-weights, and RMA values from in acute ethanol- and saline-exposed mouse PFC, VTA, and NAc for edge-weights. By the EW-dmGWAS algorithm, higher node-weights represent lower (i.e. more significant) GWAS p-values, whereas higher edge-weights represent a greater response difference of two genes between ethanol and control groups. This is calculated by taking the difference of correlations in RMA expression values of the two genes in control vs. ethanol treated BXD lines. The module score algorithm incorporated edge-and node-weights, which were each weighted to prevent bias towards representation of nodes or edges in module score calculations. Higher module scores represent higher edge- and node-weights. Genes were kept in a module if they increased the standardized module score (Sn) by 0.5%. S_n_ corresponding to a permutation-based, empirical *qFDR*<0.05 were considered significant. A significant S_n_ (i.e. more significant *qFDR* values) indicates that a module’s constituent genes are more highly associated with AD in humans, and their interactions with each other are more strongly perturbed by acute ethanol exposure in mice than randomly constructed modules of the same size.

Due to the redundancy of genes between modules, we modified the EW-dmGWAS output by iteratively merging significant modules that overlapped >80% until no modules had >80% overlap, for each brain region. Percent overlap represented the number of genes contained in both modules (for every possible pair) divided by the number of genes in the smaller module. We call the final resulting modules “mega-modules”. Standardized mega-module scores (MM-S_n_) were calculated using the algorithms employed by EW-dmGWAS. MM-S_n_ corresponding to *qFDR*<0.05 were considered significant (Fig S1). Finally, connectivity (k) and Eigen-centrality (EC) were calculated using the igraph R package for each gene in each module to identify hub genes. Nodes with EC>0.2 and in the top quartile for connectivity for a module were considered to be hub genes.

#### Overlap with ALSPAC

Genes with an ALSPAC GWAS gene-wise *p*<0.001 were considered nominally significant, and will be referred to as “ALSPAC-nominal genes” from here on out. We used linear regression to test MM-Sn’s prediction of mean ALSPAC GWAS gene-wise p-value of each mega-module. Given our hypothesis that EW-dmGWAS would identify alcohol-associated gene networks and prioritize them by association, we predicted that higher MM-S_n_’s would predict lower (i.e. more significant) mean GWAS p-values. Empirical p-values<0.017, reflecting Bonferroni correction for 3 independent tests (one per brain region): α=0.05/3, were considered to represent significant association.

Overrepresentation of ALSPAC-nominal genes within each mega-module was analyzed for those modules containing >1 such gene. For each of these mega-modules, 10,000 modules containing the same number of genes were permuted to determine significance. Empirical p-values < 0.05/n (where n = total number of mega-modules tested) were considered significant.

#### Functional enrichment analysis

To determine if mega-modules with significant overrepresentation of ALSPAC-nominal genes represented an aggregation of functionally related genes, ToppGene (https://toppgene.cchmc.org/) was used to analyze functional enrichment. Categories of biological function, molecular function, cellular component, mouse phenotype, human phenotype, pathways, and drug interaction were tested for over-representation. Significant over-representation results were defined as p<0.01 (uncorrected), n≥3 genes overlap and n≤1000 genes per functional group. Given the number of categories and gene sets tested, our discussion below was narrowed to the most relevant categories, defined as Bonferroni-corrected *p*<0.1.

## Results

Of the initial 45,037 probesets for the mouse gene expression arrays, 16,131 were associated with human-mouse homologues and had *qFDR*<0.1 for ethanol responsiveness (S-score) in at least one of the three brain regions (Fig 1). These probesets corresponded to a total of 7,730 genes and were trimmed to a single probeset per gene by filtering for the most abundant probeset as described in Methods. After removing genes that were absent from either the PPI network or the IASPSAD dataset, the final background PPI network for EW-dmGWAS analysis contained 6,050 genes (nodes) and 30,497 interactions (edges). The nodes contained 25 of the 78 IASPSAD-nominal genes and 24 of the 100 ALSPAC-nominal genes. There was no overlap between the IASPSAD and ALSPAC nominal gene sets.

### Prefrontal Cortex

For analysis using PFC expression data for edge-weights, results revealed 3,545 significant modules (*qFDR*<0.05) containing a total of 4,300 genes, with 14 ALSPAC-nominal genes and 18 IASPSAD-nominal genes. These modules were merged to form 314 mega-modules, all with significant MM-Sn. Twelve mega-modules contained at least one ALSPAC-nominal gene, and 160 contained at least one IASPSAD-nominal gene. However, MM-S_n_ did not significantly predict mean ALSPAC GWAS gene-wise p-value (*β*=−0.003, *p*=0.327, Fig 2).

**Fig 2.**
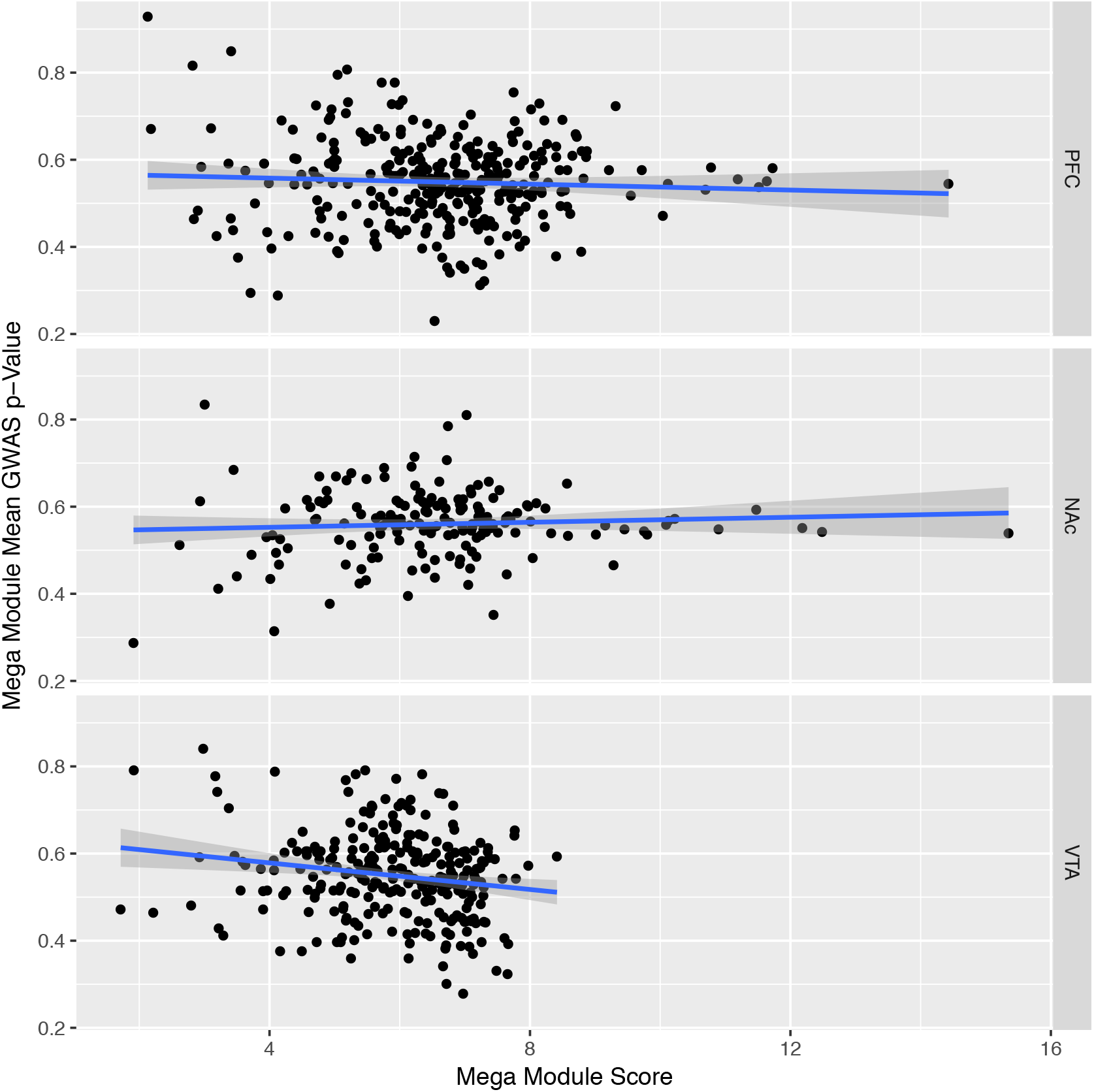
Mega Module Score v. Module Average ALSPAC GWAS p-Value. Correlation between each Mega Module’s score and average ALSPAC gene-wise GWAS p-value, for the Prefrontal Cortex (PFC) (β=−0.003, p=0.327), Nucleus Accumbens (Nac) (β=0.003, p=0.390), and Ventral Tegmental Area (VTA) (β=−0.02, p=0.003). Blue lines represent the line of best fit, estimated by linear regression, surrounded by their 95% confidence intervals (shaded gray).

Two mega-modules, Aliceblue and Cadetblue, contained multiple ALSPAC-nominal genes (Table 1). Because overrepresentation was tested for 2 mega-modules, *p*<0.025 (α=0.05/2) was considered significant. Cadetblue, was significantly overrepresented with ALSPAC-nominal genes (Table 1). Each of Cadetblue’s ALSPAC- and IASPSAD-nominal genes was connected to one of its most highly connected hub genes, *ESR1* (estrogen receptor 1; connectivity (k)=31, Eigen-centrality (EC)=1) and *ARRB2* (beta-arrestin-2; k=13, EC=0.25) (Fig 3). Although the ALSPAC-nominal gene overrepresentation was not significant for Aliceblue, it approached significance (Table 1). Further, Aliceblue had the second-highest MM-S_n_ in the PFC and contained 3 ALSPAC-nominal genes and 3 IASPSAD-nominal genes (Table 1). For these reasons, Aliceblue was carried through to functional enrichment analysis. Aliceblue’s two hub genes were *ELAVL1* ((embryonic lethal, abnormal vision)-like 1; k=165, EC=1) and *CUL3* (cullin 3; k=75, EC=0.21), which were connected to two of the three ALSPAC-nominal genes. Of these, *CPM*’s (carboxypeptidase M’s) only edge was with *ELAVL1*, and *EIF5A2’s* (eukaryotic translation initiation factor 5A2’s) only edge was with *CUL3* (Fig 3).

**Fig 3.**
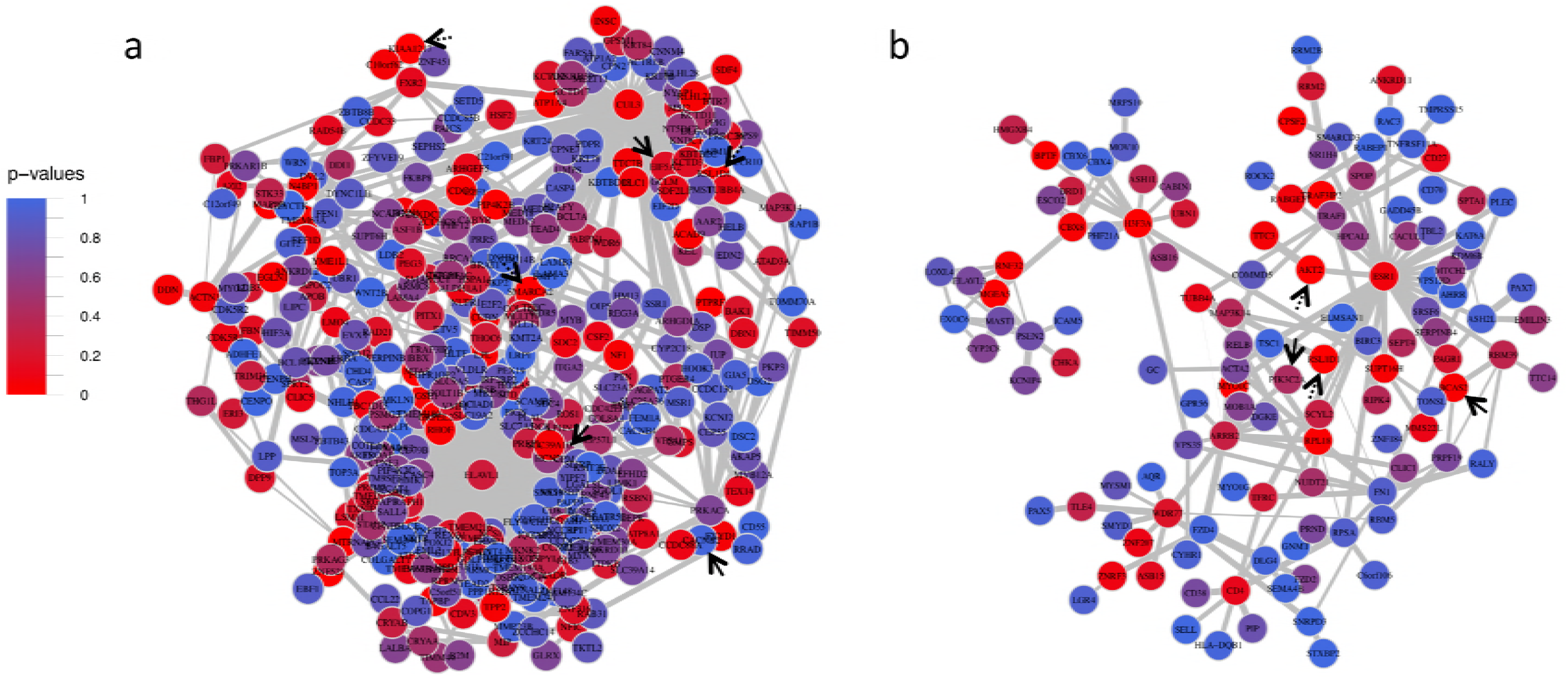
Prefrontal Cortex Mega Modules Aliceblue and Cadetblue. Prefrontal Cortex Mega Modules Cadetblue (a) and Aliceblue (b). Solid black arrows point to ALSPAC GWAS nominal genes, and dotted black arrows represent IASPSAD nominal genes. Edge-width represents strength of correlation of expression changes between treatment and control mice, and node color represents IASPSAD GWAS p-values.

**Table 1.**
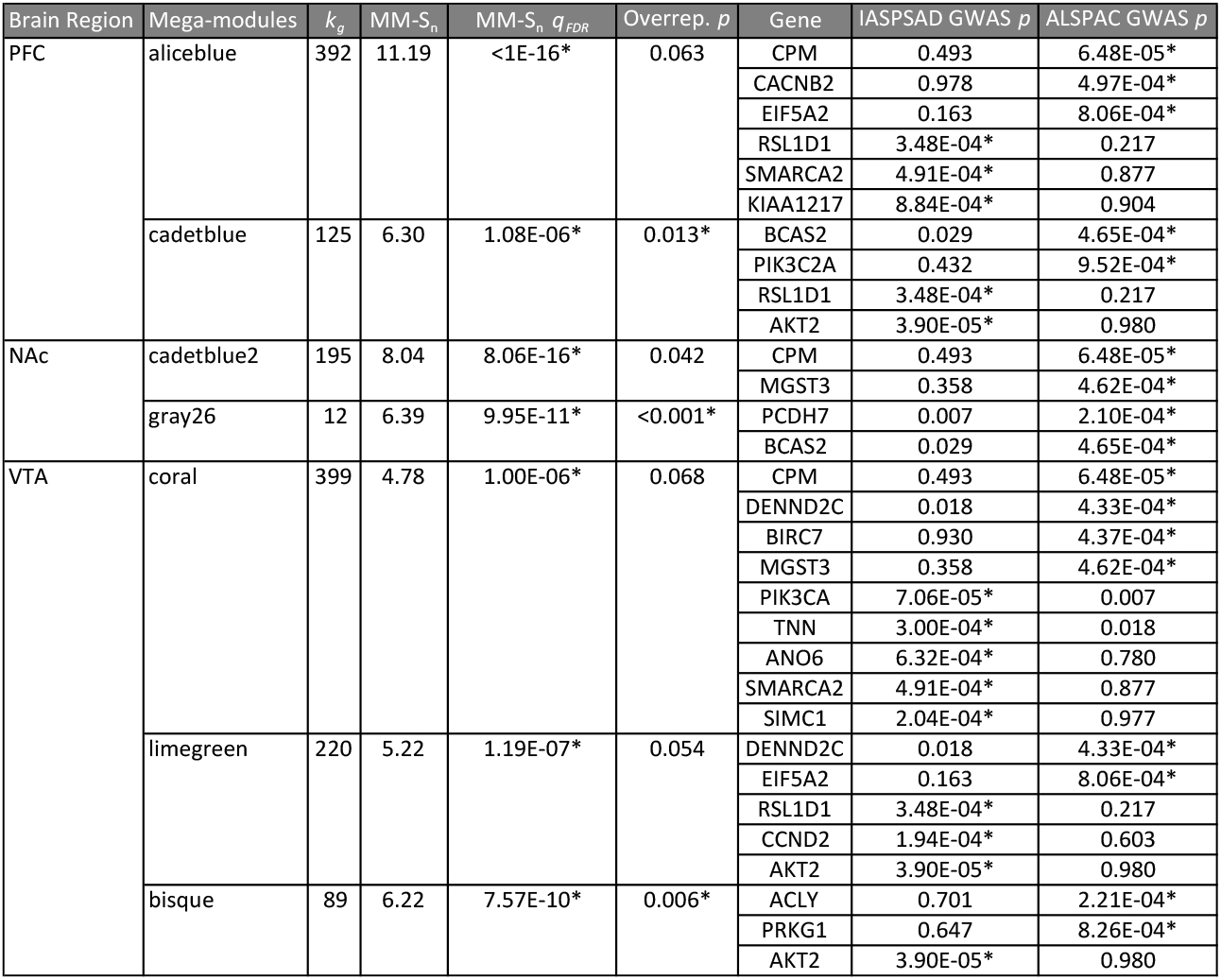
ALSPAC Nominal Gene Overrepresentation. The following characteristics are displayed for each mega-module that contained >1 ALSPAC-nominal gene: affiliated brain region; total number of constituent genes (kg); constituent ALSPAC- and IASPSAD-nominal genes; empirical p-values for ALSPAC-nominal overrepresentation (Overrep. p); MM-Sn,and the associated False Discovery Rate (MM-S_n_ qFDR). * p<0.05 for MM S_n_ and p<0.05/n for ALSPAC overrepresentation, where n=number of tests per brain region

Both Cadetblue and Aliceblue showed significant enrichment in several functional categories (Table S3). In sum, top functional enrichment categories for Aliceblue were related to actin-based movement, cardiac muscle signaling and action, increased triglyceride levels in mice, cell-cell and cell-extracellular matrix adhesion, and syndecan-2-mediated signaling. In contrast, Cadetblue’s top enrichment categories involved transcription-regulatory processes, specifically: RNA splicing, chromatin remodeling, protein alkylation and methylation, DNA replication regulation, several immune-related pathways, *NF-κβ* and Wnt signaling pathways, and reductase activity (Tables 2a-b; Table S3).

**Table 2.**
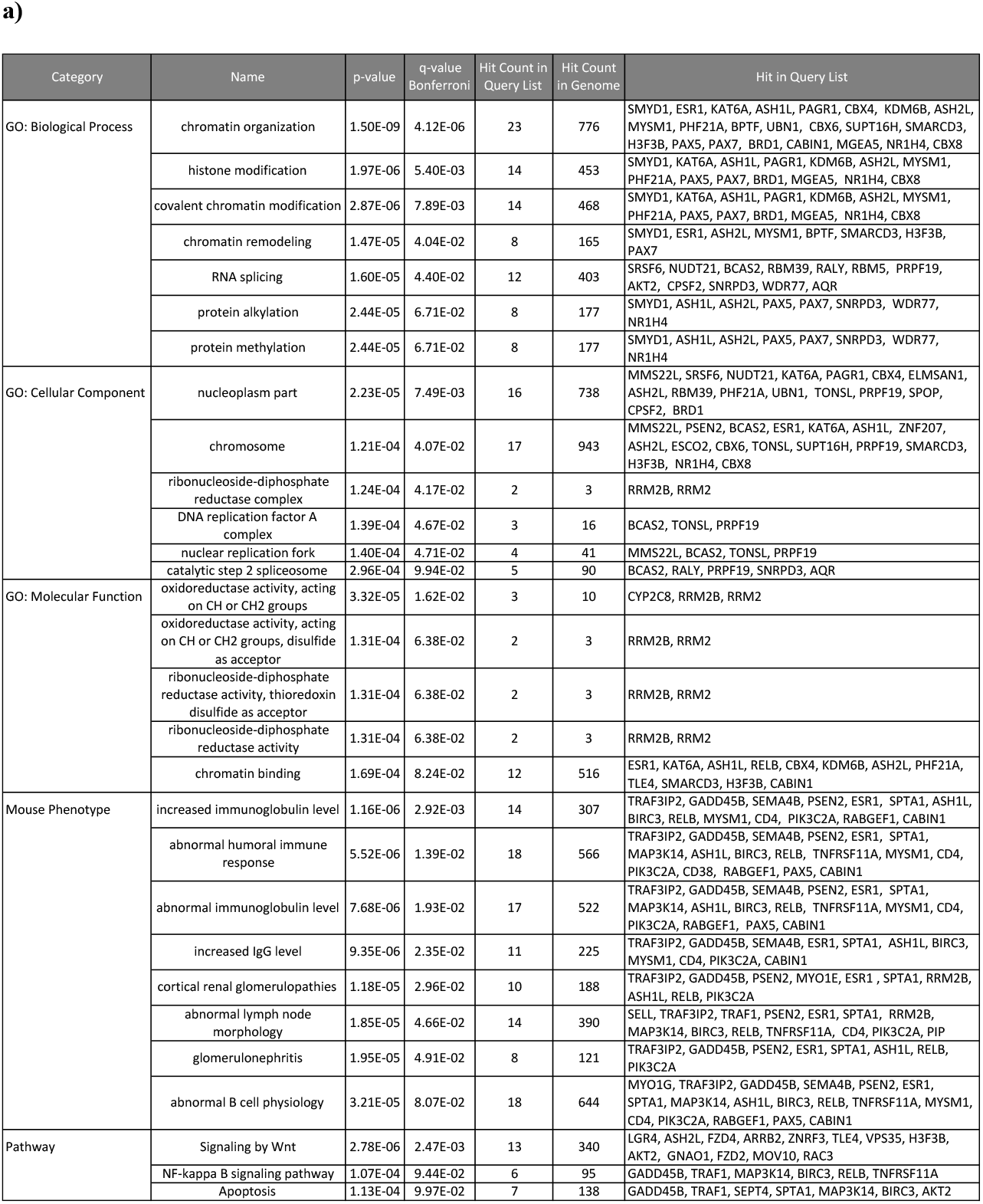
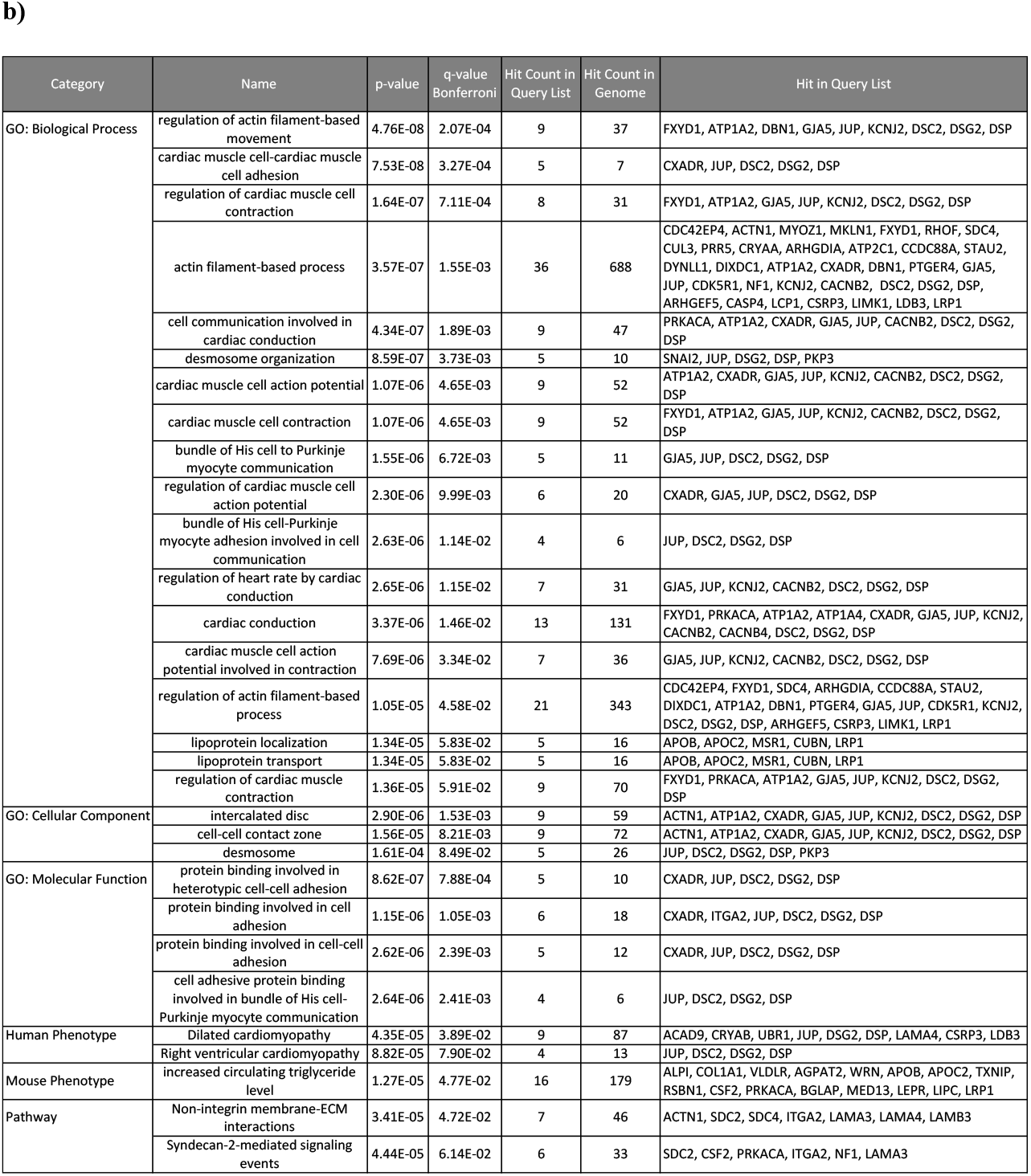
Top Gene Ontology Enrichment Results for PFC Mega Modules Cadetblue and Aliceblue. Functional enrichment results from ToppFun for Prefrontal Cortex Mega Modules Cadetblue (a) and Aliceblue (b), where Bonferroni-corrected p<0.1.

### Nucleus Accumbens

Using NAc acute ethanol expression data for edge-weights yielded 3,460 significant modules containing a total of 4,213 genes, 15 of which were ALSPAC-nominal and 16 of which were IASPSAD-nominal. After merging by content similarity, there were 171 significant mega-modules. Nineteen MM contained at least one ALSPAC-nominal gene, and 73 MM contained at least one IASPSAD-nominal gene. However, MM S_n_ did not significantly predict MM mean ALSPAC GWAS gene-wise p-value (*β*=0.003, *p*=0.390). Two MMs, Cadetblue2 and Gray26, each contained two ALSPAC-nominal genes (Table 1). Because there were 2 tests for overrepresentation, *p*<0.025 (α=0.05/2) was considered significant. Gray26, was significantly overrepresented with ALSPAC-nominal genes, and Cadetblue2 showed a trend towards overrepresentation with significance before correcting for multiple testing (Table 1).

Gray26’s most central hub gene was *HNRNPU* (heterogeneous nuclear ribonucleoprotein U; connectivity=6, Eigen-centrality=1), followed by *RBM39* (RNA binding motif protein 39; k=3, EC=0.46) and *CSNK1A1* (k=3, EC=0.37). The two ALSPAC-nominal genes *BCAS2* (breast carcinoma amplified sequence 2) and *PCDH7* (protocadherin 7), shared their only edges with *RBM39* and *HNRPNPU*, respectively (Fig 4a). As seen in the PFC’s Aliceblue, *EAVL1* was a hub gene of Cadetblue2. *ELAVL1* (k=136, EC=1) was connected to both of the ALSPAC-nominal genes, and served as the only connection for *CPM* and one of two connections for *MGST3* (microsomal glutathione S-transferase 3) (Fig 4b). Strikingly, PFC Aliceblue and NAc Cadetblue 2 showed a highly significant overlap in their gene content, with 72 overlapping genes (Table S2; p=2.2 x 10^−16^).

**Fig 4.**
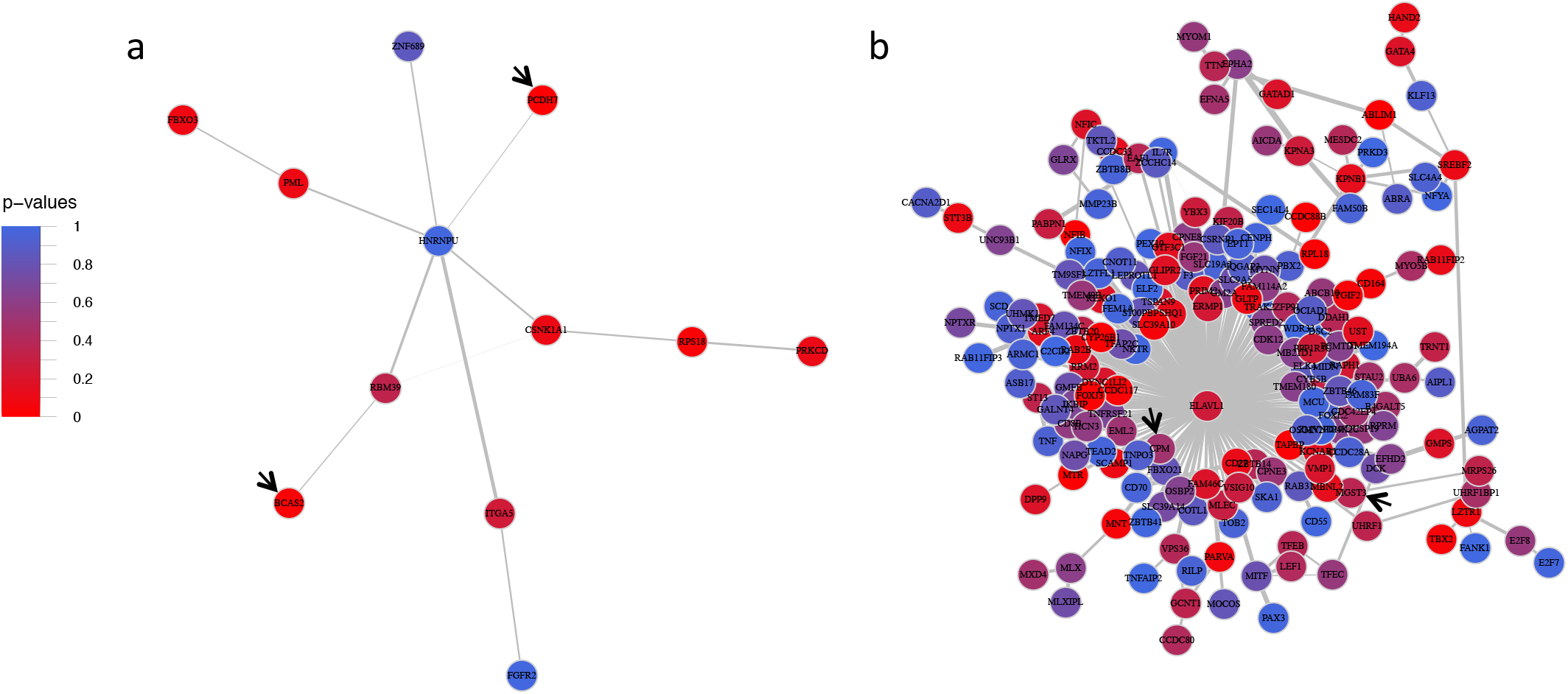
Nucleus Accumbens Mega Modules Gray26 and Cadetblue2. Nucleus Accumbens Mega Modules Gray26 (a) and Cadetblue2 (b). Solid black arrows point to ALSPAC GWAS nominal genes. These modules did not contain IASPSAD nominal genes. Edge-width represents strength of correlation of expression changes between treatment and control mice, and node color represents IASPSAD GWAS p-values.

Both Cadetblue2 and Gray26 were significantly enriched with several functional categories (Table S3). Like PFC Cadetblue, NAc Cadetblue2 was functionally enriched for gene groups related to nuclear function with transcription regulation pathways, particularly those involving RNA polymerase activity. Gray26 was most significantly enriched with genes related to functions involving: telomere maintenance, organelle organization, ribonucleoprotein complexes, and syndecan-mediated signaling (Tables 3a-b; Table S3).

**Table 3.**
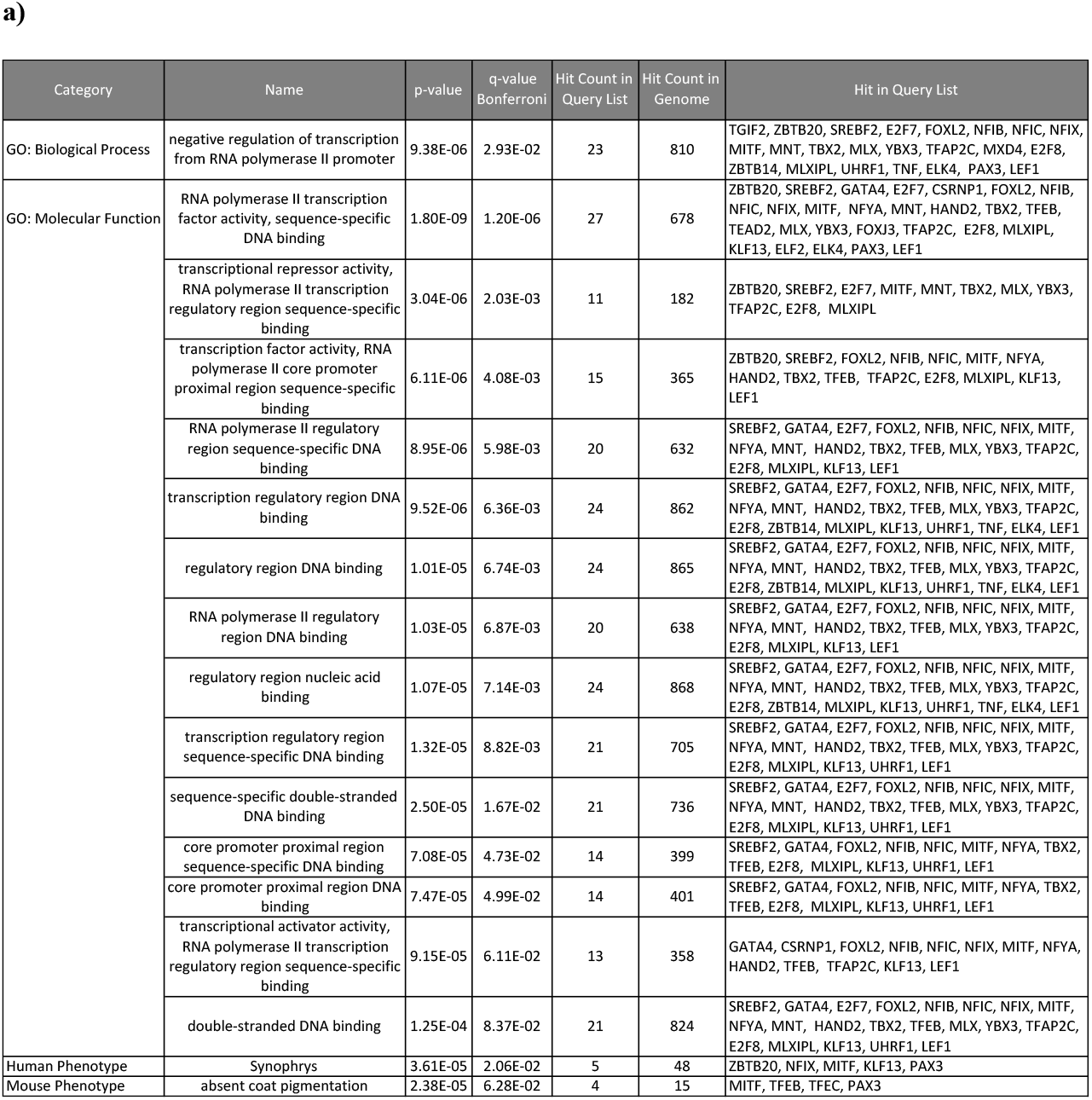
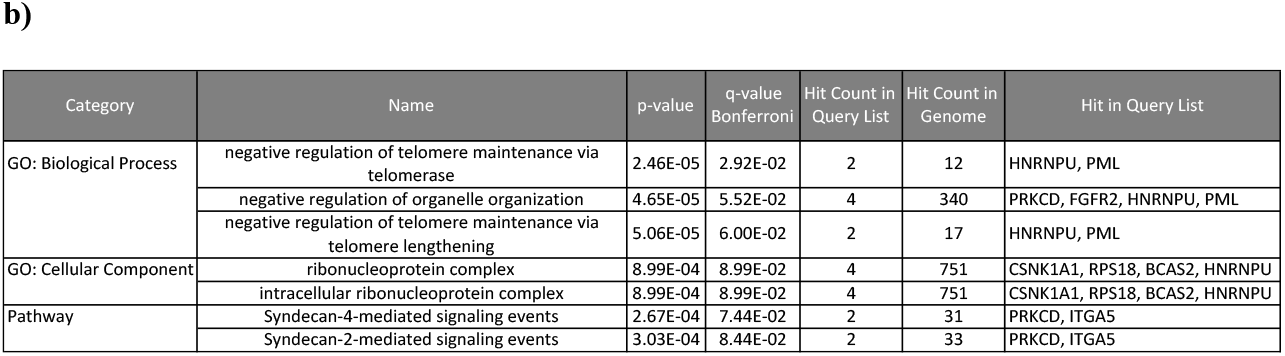
Top Gene Ontology Enrichment Results for Nucleus Accumbens Mega Modules Cadetblue2 and Gray26. Functional enrichment results from ToppFun for Nucleus Accumbens Mega Modules Cadetblue2 (a) and Gray26 (b), where Bonferroni-corrected p<0.1.

### Ventral Tegmental Area

Use of VTA control/ethanol gene expression responses for edge weighting initially resulted in 3,519 significant modules containing a total of 4,188 genes in EW-dmGWAS analysis. Merging by content similarity, resulted in 276 MMs, each with a significant MM Sn. Seventeen ALSPAC-nominal genes and 19 IASPSAD-nominal genes were spread across 25 and 156 mega-modules, respectively. Furthermore, MM-S_n_ significantly predicted mean ALSPAC GWAS gene-wise *p*-value (*β*=−0.02, *p*=0.003).

Mega-modules with the highest representation of ALSPAC-nominal genes included Coral, Limegreen, and Bisque (Table 1). Because there were 3 tests for overrepresentation, *p*<0.017 (α=0.05/3) was considered significant. Although overrepresentation of ALSPAC-nominal genes was not significant in Coral and Limegreen, it was significant in Bisque, which has the highest MM-S_n_ of the three (Table 1; Fig 5). Bisque contained four highly interconnected genes: *USP21* (ubiquitin specific peptidase 21; k=10, EC=1), *USP15* (ubiquitin specific peptidase 15; k=10, EC=0.65), *TRIM25* (tripartite motif-containing 25; k=10, EC=0.49), and *HECW2* (HECT, C2 and WW domain containing E3 ubiquitin protein ligase 2; k=12, EC=0.48). *HECW2* and *TRIM25* shared edges with this MM’s IASPSAD-nominal genes *PRKG1* (protein kinase, cGMP-dependent, type I) and *ACLY* (ATP citrate lyase), respectively. However, none of the hub genes shared an edge with Bisque’s ALSPAC nominal gene, *AKT2* (AKT serine/threonine kinase 2). Finally, Bisque had significant enrichment in several functional categories (Table S3). It was most significantly enriched with genes associated with ubiquitination, ligase and helicase activity, and eukaryotic translation elongation (Table 4; Table S3).

**Fig 5.**
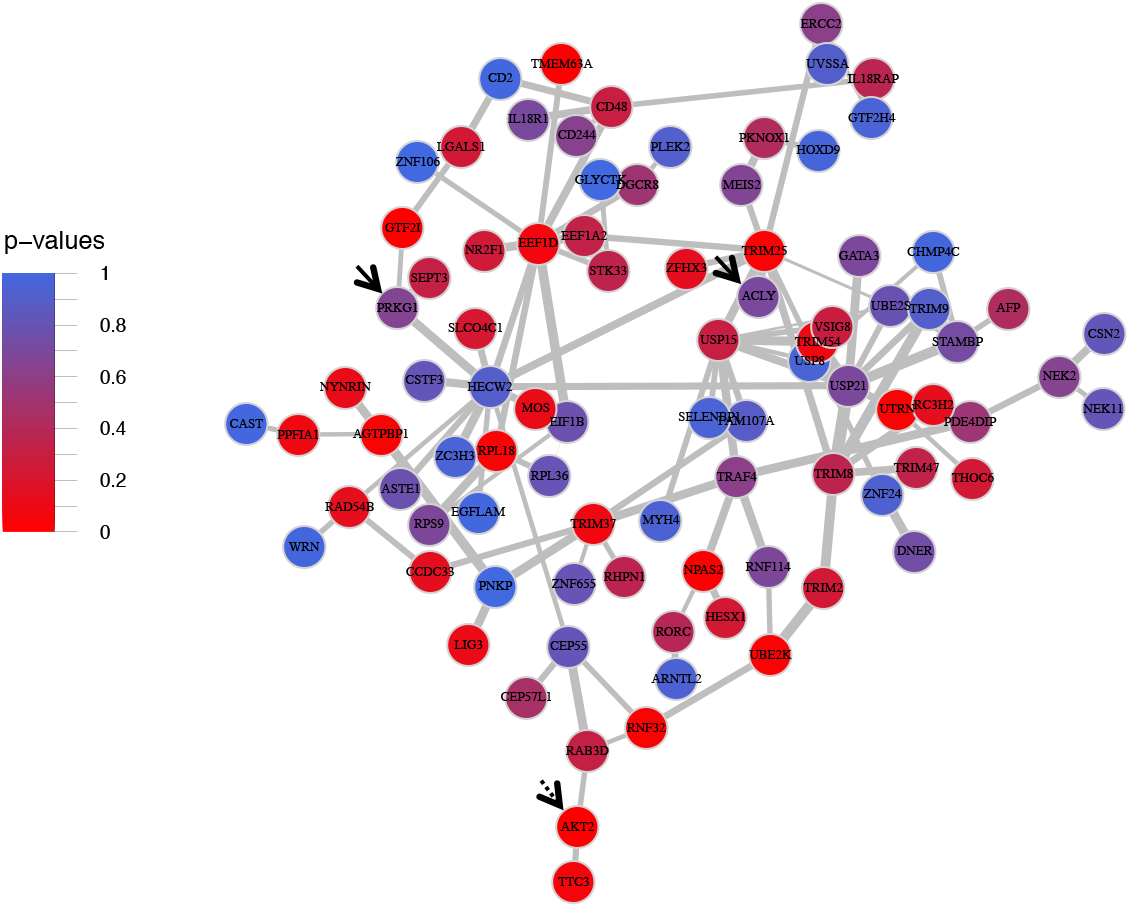
Ventral Tegmental Area Mega Module Bisque. Ventral Tegmental Area Mega Modules Bisque. Solid black arrows point to ALSPAC GWAS nominal genes, and dotted black arrows represent IASPSAD nominal genes. Edge-width represents strength of correlation of expression changes between treatment and control mice, and node color represents IASPSAD GWAS p-values.

**Table 4.**
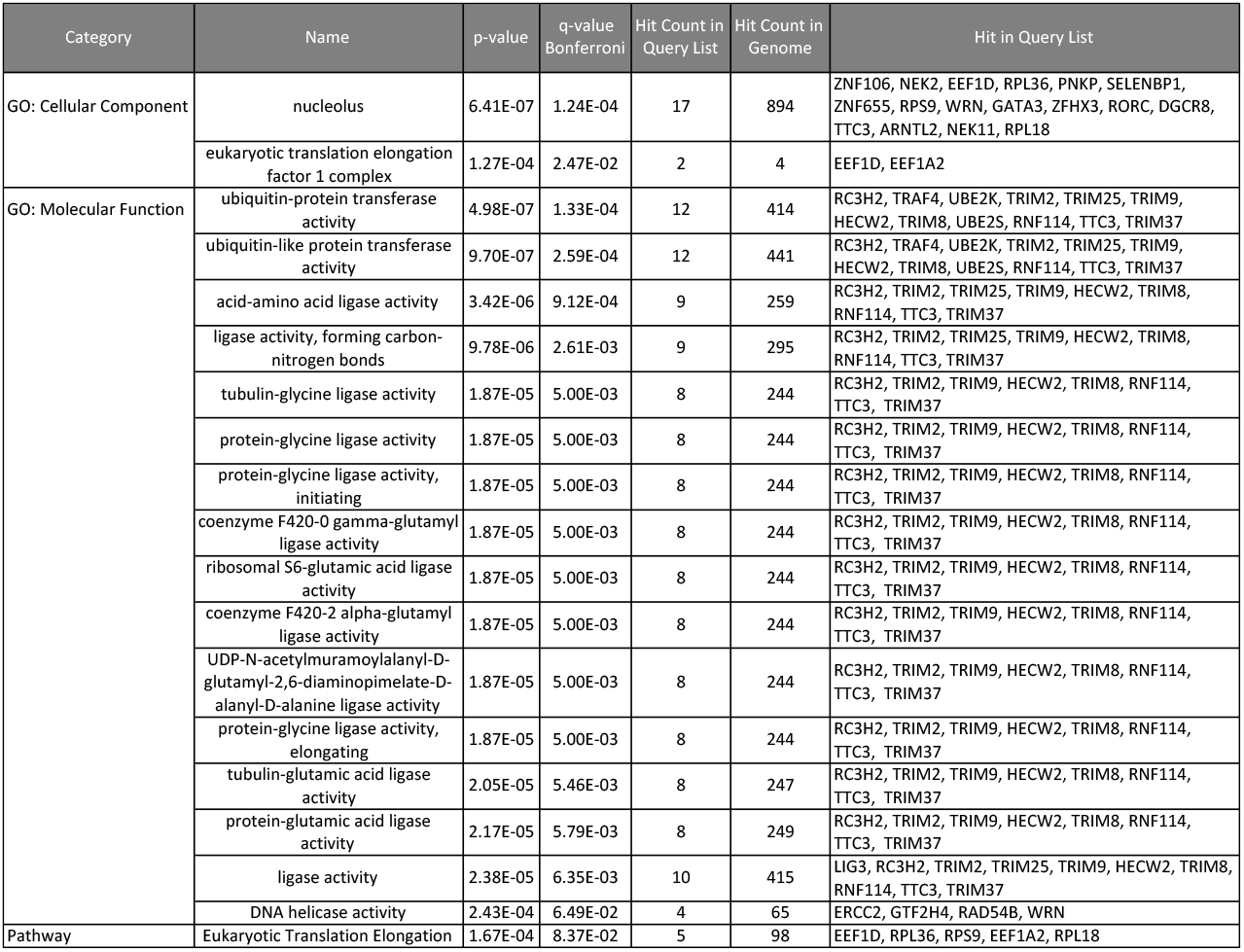
Top Gene Ontology Enrichment Results for Ventral Tegmental Area Mega Module Bisque. Functional enrichment results from ToppFun for Ventral Tegmental Area Mega Module Bisque, where Bonferroni-corrected p<0.1.

## Discussion

To our knowledge, this is the first study to directly co-analyze human GWAS with mouse brain ethanol-responsive gene expression data to identify ethanol-related gene networks relevant to AD. Unlike previous studies that have employed cross-species validation methods for specific genes or gene sets, this study analyzed human and mouse data in tandem to identify gene networks across the entire genome, using the EW-dmGWAS algorithm. This approach successfully identified significantly ethanol-regulated and AD-associated gene networks, or modules. We further improved the existing EW-dmGWAS algorithm by merging highly redundant modules to create more parsimonious mega-modules, thus decreasing complexity without sacrificing significance. Additionally, we validated these results by testing for overrepresentation with, and mega-module score prediction by, signals from an independent GWAS dataset. Overall, our findings suggest that such direct integration of model organism expression data with human protein interaction and GWAS data can productively leverage these data sources. Furthermore, we present evidence for novel, cross-validated gene networks warranting further study for mechanisms underlying AUD.

### Identification of network-level associations across GWAS datasets

One major concern with existing GWAS studies on AD had been the relative lack of replication across studies. Although some very large GWAS studies on alcohol consumption have shown replicable results [13–15], those do not account for all previously identified associations. We reasoned that our integrative gene network-querying approach might identify networks that shared signals from different GWASs on AD, even if the signals were not from the same genes across GWASs. Concordant with this hypothesis, VTA mega-module scores significantly predicted average gene-wise p-values from an independent GWAS dataset, ALSPAC (Fig 2). This suggests that ethanol-regulated gene expression networks in this brain region may be particularly sensitive to genetic variance and thus are highly relevant to mechanisms contributing to risk for AD. This is possibly attributable to the involvement of VTA dopaminergic reward pathways in the development of AD [41].

Although scores did not prioritize mega-modules with respect to ALSPAC results in PFC and NAc, individual mega-modules were overrepresented with ALSPAC signals (Table 1). The ALSPAC-overrepresented VTA and PFC mega-modules also contained nominally significant genes from the GWAS dataset used for the network analysis, IASPSAD. These results suggest that the integration of acute ethanol-related expression data from mice and human PPI can identify functional networks that associate signals from different GWAS datasets.

### Composition and structure of mega-modules

Functional composition of mega-modules varied between brain regions for the most part. For example, although Aliceblue (PFC) and Cadetblue2 (NAc) shared the hub gene *ELAVL1*, ALSPAC-nominal gene *CPM*, and had a significant overlap in their gene content, their functional enrichment results were very different (Tables 2b and 3a). These results suggest that brain regional ethanol-responsive gene expression results likely had an important impact on composition of networks, thus leveraging protein-protein interaction network information and GWAS results.

Despite such differences, the mega-modules presented in Table 1 shared certain structural similarities. Most of the IAPSAD- and ALSPAC-nominal genes in these modules shared edges with hub genes (Fig 3–5). These hub genes included: *CUL3 and ELAVL1* from PFC Aliceblue; *ESR1* from PFC Cadetblue; *ELAVL1rom* NAc Cadetblue2; *TRIM25* and *HECW2* from VTA Bisque. Further, GWAS nominally significant genes (IASPAD or ALSPAC) generally were not hub genes in the derived networks (see Fig 3–5; Table S2). This may be consistent with the general tenet that genetic variation in complex traits does not produce major alterations in cellular function, but rather modulation of cellular mechanisms for maintaining homeostasis. Hub genes may be more functionally more closely related to a given trait, but likely have such widespread influence so as to be evolutionarily resistant to genetic variation in complex traits. This is also consistent with the hypothesis that omnigenic influences are an important feature of complex traits such as AUD [42].

One hub gene was found to influence network structure in both PFC and NAc. *ELAVL1* is a broadly expressed gene that acts as a RNA-binding protein in AU-rich domains, generally localized within 3’-UTRs of mRNA. As such, *ELAVL1* has been shown to alter mRNA stability by altering binding of miRNA or other factors influencing mRNA degradation [43] and has been implicated in activity-dependent regulation of gene expression in the brain with drug abuse [44]. The large interaction space for *ELAVL1* in PFC Alice Blue and NAc Cadetblue 2 and the multiple nominal GWAS hits within these genes suggest that *ELAVL1* could have an important modulatory function on the network of genes susceptible to genetic variation in AUD.

### Functional aspects of mega-modules

This theory regarding network structure is further supported by our functional enrichment analysis, which revealed several small groups of functionally related genes within each mega-module. All of the mega-modules discussed above (Table 1) contained at least one GWAS-nominal gene in the top enrichment groups, except Cadetblue2, which still had GWAS-nominal genes in its significant enrichment groups (Table S3).

Another unifying feature across these mega-modules, except Aliceblue, was significant functional enrichment for pathways that regulate gene expression. Specifically, these pathways were related to chromatin organization, RNA splicing, and translation- and transcription-related processes (Table S3). This is not surprising, as alterations in gene expression have long been proposed as a mechanism underlying long-term neuroplasticity resulting in ethanol-dependent behavioral changes, and eventually dependence [45].

In contrast, the largest functional enrichment groups unique to Aliceblue were related to actin-based filaments and cardiac function (Table 2). Actin not only provides cytoskeletal structure to neurons, but also functions in dendritic remodeling in neuronal plasticity, which likely contributes to AD development [46, 47]. Aliceblue was also significantly enriched for the syndecan-2 signaling pathway, and contained the *SDC2* gene itself, which functions in dendritic structural changes together with F-actin [48]. Additionally, the most significant enrichment group unique to Cadetblue was the Wnt signaling pathway, which also regulates actin function [49, 50]. Of note, a prior study has shown that *ARRB2* (a Cadetblue hub gene and member of Wnt signaling pathway) knockout rats display significantly decreased levels of voluntary ethanol consumption and psychomotor stimulation in response to ethanol [51]. These findings highlight the potential importance of postsynaptic actin-related signaling and dendritic plasticity in PFC gene networks responding to acute ethanol and contributing to genetic risk for AD.

Finally, although the NAc Cadetblue2 mega-module was highly enriched for functions related to transcriptional regulation, it also contained the gene *FGF21* within its interaction space (Table S2 and Fig 4b). FGF21 is a member of the fibroblast growth factor gene family and is a macronutrient responsive gene largely expressed in liver. Importantly FGF21 has been shown to be released from the liver by ethanol consumption and negatively regulates ethanol consumption by interaction with brain FGF-receptor/beta-Klotho complexes. Beta-Klotho, a product of the *KLB* gene, is an obligate partner of the FGF receptor and has recently been shown to have a highly significant association with alcohol consumption in recent very large GWAS studies [14, 15]. Although the role of *FGF21* and *KLB* in AD are not currently known, the association of *FGF21* with the Cadetblue2 mega-module, containing nominally responsive genes from AD GWAS studies, is a possible additional validation of the utility of our studies integrating protein-protein interaction information (tissue non-specific), AD GWAS (tissue non-specific) and brain ethanol-responsive gene expression.

### Potential weaknesses and future studies

The studies presented here provide evidence for the utility of integrating genomic expression data with protein-protein interaction networks and GWAS data in order to gain a better understanding of the genetic architecture of complex traits, such as AD. Our analysis also generated several testable hypotheses regarding gene networks and signaling mechanisms related to ethanol action and genetic burden for AD. However, these studies utilized acute ethanol-related expression data in attempting to identify mechanisms of AD, a chronic ethanol exposure disease. Use of a chronic exposure model could provide for a more robust integration of the expression data and GWAS signals. However, we feel the current study is valid, since acute responses to ethanol have been repeatedly shown to be a heritable risk factor for AD [52–54].

Another potential shortcoming for this work regards the limited size of the GWAS studies utilized and differences in phenotypic assessment. The IASPSAD study was based on AD diagnosis, whereas ALSPAC was based on a symptom factor score. Had we used larger GWAS studies based on the same assessment criteria, it is possible that greater overlap of GWAS signals within mega-modules would have been observed. Recent large GWAS studies on ethanol have, to date, generally concerned measures of ethanol consumption, rather than a diagnosis of alcohol dependence per se [14, 15]. For this reason, we focused this initial effort on GWAS studies concerned with alcohol dependence. However, using the IASPAD and ALSPAC studies allowed us to identify gene networks that are robust across both the severe end of the phenotypic spectrum (i.e. diagnosable AD), and for symptoms at the sub-diagnostic level.

Overall, this analysis successfully identified novel ethanol-responsive, AD-associated, functionally enriched gene expression networks in the brain that likely play a role in the developmental pathway from first ethanol exposure to AD, especially in the VTA. This is the first analysis to identify such networks by directly co-analyzing gene expression data, protein-protein interaction data, and GWAS summary statistics. The identified modules provided insight into common pathways between differing signals from independent, largely underpowered, yet deeply phenotyped GWAS datasets. This supports the conjecture that the integration of different GWAS results at a gene network level, rather than simply looking for replication of individual gene signals, could make use of previously underpowered datasets and identify common genetic mechanisms relevant to AD. Future expansion of such approaches to include larger GWAS datasets and chronic ethanol expression studies, together with validation of key targets by gene targeting in animals models, may provide both novel insight for the neurobiology of AD and the development of improved therapeutic approaches.

## Acknowledgements

The authors wish to thank members of the Miles laboratory and the Virginia Institute for Psychiatric and Behavioral Genetics for their helpful comments and suggestions during the course of these studies. Additionally, we thank Dr. Zhongming Zhou at the University of Texas Health Sciences Center for suggestions regarding the utilization of EW-dmGWAS. None of the authors had any conflicts of financial interest in the performance of these studies.

## Supporting information

**S1 Fig. Analytical Pipeline of Steps Following EW-dmGWAS**. Empirical p-values were calculated from standardized module scores based on a Z-distribution. The original EW-dmGWAS module score, permutation, and score standardization algorithms were used to calculate the respective Mega Modules parameters. Modules were considered to have >80% overlap if >80% of the genes in the smaller module was contained in the larger module. False Discovery Rates were calculated based on the Benjamini-Hochberg algorithm, using the “stats” package in R. Intramodular connectivity was defined as the number of edges (i.e. connections) attached to that node (i.e. gene). Eigen-Centrality was calculated using the “igraph” package in R.

**S1 Table. Brain Region-Specific S-score Values**. One table per brain region, containing each of the following values: RMA values and S-scores from the maximally expressed probeset per gene, for each BXD strain; the associated probeset IDs, human gene symbols, and mouse gene symbols; and the Fisher’s combined False Discovery Rate (q-value) for each probeset.

**S2 Table. Mega Module Characteristics**. One table per brain region, containing each of the following characteristics, for all significant Mega Modules: name; constituent genes; ALPSAC and IASPSAD p-values for each gene; Mega Module score (S_n_), p-value (S_n__p), and False Discovery Rate (S_n__qFDR); and intramodular eigencentrality and connectivity. Significance values < 10^−16^ are rounded to 0.

**S3 Table. Mega Module Gene Ontology Enrichment**. One table for each ALSPAC-overrepresented Mega Module, containing ToppFun output for gene ontology enrichment groups with *p*<0.01 and minimum group size of 3 genes and maximum size of 1,000 genes, for the following categories: Biological Process, Cellular Component, Molecular Function, Human Phenotype, Mouse Phenotype, and Pathways.

